# Comparative Proteomics Reveals Characteristic Proteins on Praziquantel-resistance in *Schistosoma mansoni*

**DOI:** 10.1101/314724

**Authors:** António Pinto-Almeida, Tiago Mendes, Pedro Ferreira, Silvana Belo, Fernanda de Freitas Anibal, Silmara Marques Allegretti, Emanuel Carrilho, Ana Afonso

**Affiliations:** Graduate Program in Areas of Basic and Applied Biology, Instituto de Ciências Biomédicas Abel Salazar, Universidade do Porto, Porto, Portugal; Global Health and Tropical Medicine, GHTM, UEI Medical Parasitology, Instituto de Higiene e Medicina Tropical, IHMT, Universidade Nova de Lisboa, UNL, Lisbon, Portugal; São Carlos Institute of Chemistry, Universidade de São Paulo, São Carlos, SP, Brazil; Institute of Biology, Animal Biology Department, Universidade Estadual de Campinas, SP, Brazil; Laboratory of Parasitology, Department of Morphology and Pathology, Universidade Federal de São Carlos, São Carlos, SP, Brazil; Faculty of Engineering and Marine Sciences, University of Cape Verde, Campus of Ribeira de Julião, Mindelo, São Vicente, Cape Verde

**Keywords:** *Schistosoma mansoni*, Praziquantel, Proteomics, Susceptibility, Resistance, Mass spectrometry

## Abstract

The extensive use of Praziquantel (PZQ), the only drug available to treat schistosomiasis, has brought concern about the emergence of PZQ-resistance/tolerance by *Schistosoma* spp., thus reaffirming an urge for the development of new treatment alternatives. Therefore, it is imperative and urgent to study this phenomenon trying to understand what is involved in its occurrence. Studies of *Schistosoma* spp. genome, transcriptome and proteome are crucial to better understand this situation. By stepwise drug pressure from a fully susceptible parasite strain, our group selected a *S. mansoni* variant strain stably resistant to PZQ and isogenic to its fully susceptible parental counterpart, except for the genetic determinants of PZQ-resistance phenotype. Based on this, the objective of this study was to compare the proteomes of both strains, identifying proteins from male and female adult worms of PZQ-resistant and PZQ-susceptible strains, exposed and not exposed to PZQ, which were separated by high-resolution two-dimensional electrophoresis and sequenced by high throughput LC-MS/MS. Likewise, this work is extremely relevant since for the first time the proteome of a *S. mansoni* PZQ-resistant strain is studied and compared to the proteome of the respective *S. mansoni* PZQ-susceptible strain. This study identified 60 *S. mansoni* proteins, some of which differentially expressed in either strain, which may putatively be involved in the PZQ-resistance phenomenon. This information represents substantial progress towards deciphering the worm proteome. Furthermore, these data may constitute an informative source for further investigations into PZQ-resistance and increase the possibility of identifying proteins related to this condition, possibly contributing to avoid or decrease the likelihood of development and spread of PZQ-resistance. This is an innovative study that opens doors to PZQ-resistance surveys, contributing to discover a solution to PZQ-resistance problem, as suggests new potential targets for study.

## 1. Introduction

Parasites from the genus *Schistosoma* are water-borne parasites that are the causative agents of schistosomiasis, one of the most important infectious parasitic diseases, being endemic in 76 countries with more than 97% of infected people living in sub-Saharan Africa. Schistosomes infect an estimated 249 million people worldwide with an additional 732 million people at risk of infection [Steinmann et al., 2006; Kamel et al., 2011; World Health Organization (WHO), 2013]. One of the major species acting as causative agent of schistosomiasis is *Schistosoma mansoni*, a mammalian parasite endemic in Africa and parts of South America (Crompton, 1999; Chitsulo et al., 2000).

Despite many efforts to control the transmission of this disease (Doenhoff et al., 2000; Fenwick et al. 2003; Hagan et al., 2004), essentially after the introduction of a chemotherapeutic treatment in 1980s, schistosomiasis is still highly prevalent (King, 2009). Its control is based on Praziquantel (PZQ) the only drug available for chemotherapy (Cioli and Pica-Mattoccia, 2003). Treatment with PZQ is effective and inexpensive (Cioli and Pica-Mattoccia, 2003), but frequent schistosome reinfection occurs in endemic areas and may cause irreversible damages to the liver, kidneys, or urinary tract (Knudsen et al., et al., 2005). Because of its high prevalence, schistosomiasis has earned a Category II disease, ranking next to malaria, for importance as a target tropical disease by the World Health Organization Special Program for Research and Training in Tropical Diseases (Knudsen et al., et al., 2005).

Although the impact of schistosomiasis could be dramatically reduced by improvement in education and sanitation for humans and elimination of the intermediate host snails, such methods are not sufficient to control or eradicate this parasitosis. In the absence of vaccines, the control of this disease relies on chemotherapy to ease symptoms and reduce transmission. The increasing reliance on mass PZQ administration programs has exerted selective pressure on parasite population and PZQ-resistance/loss of susceptibility is being described (Wang et al., 2012). With no alternative drugs or vaccines, the fight against schistosomiasis could become a huge battle (James et al., 2009).

Identification of proteins is very important for understanding how schistosomes regulate host immune systems to establish chronic infections and also elucidate other aspects of parasite-host interaction (Knudsen et al., et al., 2005). Furthermore, a comprehensive deciphering of the schistosome genome, transcriptome, and proteome has become increasingly central for understanding the complex parasite-host interplay (Hu et al., 2004; Verjovski-Almeida et al., 2004). Therefore, such information can be expected to facilitate the discovery of vaccines and new therapeutic drug targets, as well as new diagnostic reagents for schistosomiasis control (Hu et al., 2004; Verjovski-Almeida et al., 2004; Knudsen et al., et al., 2005), and may aid the development of protein probes for selective and sensitive diagnosis of schistosomiasis (Liu et al., 2009).

Proteomics approaches encompass the most efficient and powerful tools for identification of protein complexes (Clauser et al., 1999; Perkins et al., 1999; Groth et al., 2004) and have been widely used to decipher the proteome of many trematodes (Al-sherbiny et al., 2003; Smyth et al., 2003; Kumagai et al., 2005; van Balkom et al., 2005; DeMarco and Verjovski-Almeida, 2009). For *Schistosoma* spp., the proteome has been studied in many developmental life stages, including schistosomula (Harrop et al., 1999; Curwen et al., 2004), cercariae (Curwen et al., 2004; Dvorák et al., 2008; Hansell et al., 2008), egg (Curwen et al., 2004; Cass et al., 2007) and adult worm (Curwen, 2004; Braschi and Wilson, 2006; Liu et al., 2009; Ferreira et al., 2014; Ludolf et al., 2014). But to our knowledge, the *S. mansoni* PZQ-resistant strain proteome has not been yet reported and a schistosomiasis mansoni coherent screening for proteins related to PZQ-resistance is still necessary. Understanding the development of PZQ-resistance in *S. mansoni* is crucial to prolong the efficacy of the current drug and develop markers for monitoring drug resistance. It would also be beneficial in the design of new chemotherapeutic agents to overcome or prevent resistance and the identification of new drug targets.

Our research group recently developed a resistant strain of *S. mansoni* that tolerates up to 1,200 mg PZQ/kg of mouse body weight. This *S. mansoni* variant strain was selected from a fully susceptible parasite strain, by stepwise drug pressure, and is isogenic, except for the genetic determinants of PZQ-resistance phenotypes, and significantly different of the counterpart *S. mansoni* susceptible strain (Pinto-Almeida et al., 2015). As such, this *S. mansoni* PZQ-resistant strain represents a distinct and valuable model for the study of PZQ-resistance.

The present study intended to analyze, for the first time, the proteome of *S. mansoni* PZQ-resistant adult worms and compare it with its parental fully PZQ-susceptible strain, using a high throughput LC-MS/MS identification. Therefore, this study could possibly represent a substantial progress toward deciphering the worm proteome, and may constitute an informative source for further investigations into PZQ-resistance, increasing the possibility of avoid or decrease the likelihood of development and spread of PZQ-resistance.

## 2. Material and methods

### 2.1 Parasite samples

Two different parasite isolates were used in this study, a *S. mansoni* BH PZQ-susceptible strain (SS) from Belo Horizonte, Minas Gerais, Brazil, and a stable PZQ-resistant strain (RS) (IHMT/UNL) obtained from the same BH strain as described by (Pinto-Almeida et al., 2015). These two parasite strains are routinely kept in their intermediate host *Biomphalaria glabrata* snails at IHMT/UNL.

*Mus musculus* CD1 line male mice was chosen as the animal model for *S. mansoni* infection, because it is a good host for this parasite mimicking the *S. mansoni* human infection (Katz and Coelho, 2008). Mice infection occurred by natural transdermal penetration of cercariae, by exposing mice tails to about 100 cercariae of *S. mansoni* each.

Eight to ten-weeks adult worms were recovered by perfusion of the hepatic portal system and mesenteric veins, according to (Lewis, 1998), and washed twice in RPMI-1640 medium (Sigma-Aldrich), to remove contaminating hair and blood clots.

It was analyzed males and females in separate, not exposed and exposed to PZQ, for RS and SS. Regarding to the groups of adult worms exposed to PZQ (EPZQ), after collecting, the parasites were transferred to 24-well culture plates containing RPMI-1640 culture medium, 200 mM L-glutamine, 10 mM HEPES, 24 mM NaHCO_3_, 10,000 UI of penicillin and 10 mg/mL of streptomycin, from Sigma-Aldrich, pH 7 and supplemented with 15% fetal bovine serum. Adult worms of the parasite were added on each well for each studied group for PZQ treatment: 1) PZQ-susceptible male (SM); 2) PZQ-susceptible female (SF); 3) PZQ-resistant male (RM), and 4) PZQ-resistant female (RF). Adult parasites were treated in culture, with 0.3 μM of PZQ during 24 hours and then washed twice with saline solution to clean any traces of culture medium and stored in Trizol (Invitrogen) at -80 ºC, for posterior protein extraction. For the groups of adult worms not exposed to PZQ (NEPZQ), worms were kept in RPMI-1640 medium with no addition of drug, then washed twice with saline solution to clean any traces of culture medium and also stored in Trizol (Invitrogen) at −80 ºC, for posterior protein extraction. Accordingly, the experimental set up consisted of eight sample groups, four for parasites not exposed to PZQ (RM-NEPZQ, RF-NEPZQ, SM-NEPZQ and SF-NEPZQ) and four for parasites exposed to PZQ (RM-EPZQ, RF-EPZQ, SM-EPZQ and SF-EPZQ).

### 2.2 Preparation of protein extracts

*Schistosoma mansoni* adult worm protein extracts were obtained using Trizol (Invitrogen) protocol, according to the manufacturer’s instructions. Briefly, the parasites were lysed and homogenized directly in Trizol reagent at room temperature. The homogenized samples were incubated at room temperature to permit complete dissociation of the nucleoprotein complex. After homogenization, we proceeded to separation phase, adding chloroform and centrifugation of samples. The aqueous phase was removed and the interphase and organic phenol-chloroform phase was used for protein isolation procedure. Next, isopropanol precipitation was performed and the pellet was solubilized in SBI buffer [7 M urea, 2 M thiourea, 15 mM 1,2-diheptanoyl-snglycero-3-phosphatidylcholine (DHPC), 0.5% triton X-100, 20 mM dithiotheitol (DTT) (Sigma) and Complete Mini Protease Inhibitor Cocktail Tablets], according to (Babu et al., 2004), and stored at −80 ºC until use.

Protein concentration in protein extracts was measured by Bradford assay (Bradford, 1976) and the quality of the extract was verified in 12% uniform SDS-PAGE gels.

### 2.3 Two-dimensional electrophoresis

Each experiments with two-dimensional electrophoresis (2-DE) gels were performed in triplicate with 240 μg of protein, for each group. To prepare samples for 2-DE, protein samples in the mentioned concentration were diluted in rehydration solution containing 7 M urea, 2 M thiourea, 4% CHAPS (3-[3-(cholamidopropyl)dimethylammonio]-1-propanesulfonate), 0.5% IPG buffer, 1% DTT, and 0.002% bromophenol blue (Sigma). The rehydration was carried out passively overnight during 12 h in a 13 cm, pH 3-10 strip (Immobiline Drystrips, GE Healthcare). The strips were then applied on an Ettan IPGphor 3 (GE Healthcare) system, for protein separation by isoelectric focusing (IEF) following a typical IEF protocol, which involved three focusing steps at a constant 50 μA/strip: 3 h gradient to 3,500 V, 3 h at 3,500 V and finally 64,000 V h.

After focusing, the strips containing proteins were reduced in an equilibration solution (50 M Tris HCl, pH 8.8, 6 M urea, 20% glycerol, 2% SDS) containing 2% DTT, and then alkylated in the same solution containing 2.5% iodoacetamide (IAA) (Sigma). The Immobilized pH gradient (IPG) strips and molecular weight standards were then transferred to the top of 12% uniform SDS-PAGE gels and sealed with 0.5% agarose. The second dimension was carried out using a Protein Plus Dodeca cell system (Bio-Rad) under an initial current of 15 mA/gel for 15 min, followed by increasing the current to 50 mA/gel until the end of the run (the dye front reached the bottom of the gel).

For 2-DE experiments, at least three replica of two dimensional polyacrylamide gel electrophoresis were performed for each group, confirming the reproducibility of the experimental procedure. Gels were fixed in 40% methanol/10% acetic acid solution and stained with Coomassie brilliant blue R-350 (GE Healthcare). The spots were normalized and evaluated by the software ImageMaster 2D Platinum 7.0 (GE Healthcare).

### 2.4 In-gel digestion and peptide preparation for mass spectrometry analysis

The selected protein spots from two biological samples for each group were manually excised, distained, reduced, alkylated and digested in gel with trypsin (Promega) from the corresponding 2- DE gel for mass spectrometry (MS) identification. First, spots were washed in Milli-Q water, and then distained in a distaining solution containing 50% methanol/2.5% acetic acid in purified water for 2 h at room temperature. This step was repeated until clear of blue stain. The gel fragments were incubated in 100% acetonitrile (ACN) with occasional vortexing, until gel pieces became white and shrank. Then, the solution was removed and spots were completely dried, and ready for digestion. The in-gel digestion with trypsin-modified sequencing-grade reagents (Promega) was done according to (Shevchenko et al., 2006). Briefly, protein digestion was conducted at 37 ºC overnight. After the incubation, the supernatant was transferred to a clean tube and 30 μL of 5% formic acid (FA)/60% ACN were added to gel spots for the extraction of tryptic peptides. This procedure was performed 2 × 30 min under constant agitation. The supernatant was pooled to the respective tube containing the initial peptide solution. This solution was dried in a SpeedVac (Thermo Scientific) and the peptides were re-suspended in 8 μL of 0.1% FA. The peptides were desalted in reverse phase micro-columns ZipTip C18 (Millipore), according to manufacturer’s instructions. Peptides were dried again and re-suspended in 50% ACN/ 0.1% trifluoracetic acid (TFA) solution.

### 2.5 Peptide analysis by LC-MS/MS and protein identification

The digested peptides were analyzed by LC-MS/MS using a nano-LC system (EASY-nLC II, Thermo Scientific), coupled online to a hybrid mass spectrometer ion trap linear-Orbitrap (LTQ Orbitrap Velos, Thermo Scientific) using an ion nanospray source, namely, Nano-Flex II nanospray (Thermo Scientific). The samples were injected (10 μL/min, 4 min) in a pre-column (C18, 100 μm DI × 2 cm, Thermo Scientific), and then eluted under flow of 300 nL/min using an elution gradient to a C18 column (10 cm × 75 μm DI, 3 μm, 120 Å, Thermo Scientific). All LC-MS/MS data (in raw format) were acquired using XCalibur software, version 2.0.7 (Thermo Fisher Scientific) and converted in mgf files using MassMatrix MS Data File Conversion version 3.9. The analyzes were performed in scan mode in the range of 400-2000 m/z; positive mode; capillary voltage of 4500 V; nebulizer to 8.0 psi; drying gas at a flow of 5.0 L/min and at 220 ºC of evaporation temperature of the spray.

The list of peptide and fragment mass values generated by the mass spectrometer for each spot were submitted to a MS/MS ion search using the Mascot 2.0 online search engine (Matrix-Science) to search the LC-MS/MS data against the NCBInr database *Schistosoma_mansoni*_NCBI_112014, November 2014. Mascot software was set with two tryptic missed cleavages, a peptide ion mass tolerance of 10 ppm, a fragment ion mass tolerance of 0.02 Da, a peptide charge of 2+ and 3+, a variable modifications of methionine (oxidation), and a fixed modifications of cysteine (carbamidomethylation). During the analysis, our samples were checked against a contaminant database supplied by the Global Proteome Machine. All validated proteins had at least two independent spectra, with greater than 95.0% probability estimated by the Peptide Prophet algorithm of being present in the *S. mansoni* database as at least one unique peptide. To avoid random matches, only ions with individual score above of the indicated by the Mascot to identity or extensive homology (p<0.05) were considered for protein identification. However, when the Mascot score was not significant, but the percentage coverage and root mean squared error (RMSE) were in the same range as those of proteins with a significant match, proteins were deemed identified if additional parameters, such as its calculated p*I* and *M*w, were in agreement with those observed for the actual gel spot and the species matched was *S. mansoni*, according to [26].

The molecular function and biological process were assigned for the proteins identified according to information obtained from the Gene Ontology (GO) database (Huntley et al., 2015). The exact annotation for each protein was used in most cases. However, the catalytic activity category was used for all proteins with molecular function associated with (GTPase, hydrolase, isomerase, kinase, ligase, lyase, oxidoreductase, transcription and transferase activities). Binding category was used for all types of ligand identified (actin, ATP, calcium, GTP, magnesium ion, metal ion, protein domain specific and nucleotide bindings). Furthermore, there was other molecular function categories classified such as chaperone, motor, regulation of muscle contraction, structural and transport. The proteins that had no associated known function were classified as “unknown”.

### 2.6 Ethics statement

This research project was reviewed and approved by the Ethics Committee and Animal Welfare, Faculty of Veterinary Medicine, UTL (Ref. 0421/2013). Animals were maintained and handled in accordance with National and European legislation (DL 276/2001 and DL 314/2003; 2010/63/EU adopted on 22^nd^ September 2010), with regard to the protection and animal welfare, and all procedures were performed according to National and European legislation. The anesthetics and other techniques were used to reduce the pain and adverse effect of animal.

## 3. Results

### 3.1 2-DE separation of proteins from S*. mansoni* PZQ-resistant and PZQ-susceptible adult worms

As said in the Material and methods section, in total, eight protein extracts were analyzed: four from parasites not exposed to PZQ (NEPZQ) [resistant males and females (assigned as RM-NEPZQ and RF-NEPZQ, respectively), and susceptible males and females (assigned as SM-NEPZQ and SF-NEPZQ, respectively)] and another four samples from parasites exposed to PZQ (EPZQ) [resistant males and females (assigned as RM-EPZQ and RF-EPZQ, respectively), and susceptible males and females (assigned as SM-EPZQ and SF-EPZQ, respectively)]. All protein extracts presented high purity and good quality for posterior 2-DE and mass spectrometry (MS) analysis (Figure 1).

**Figure 1.**
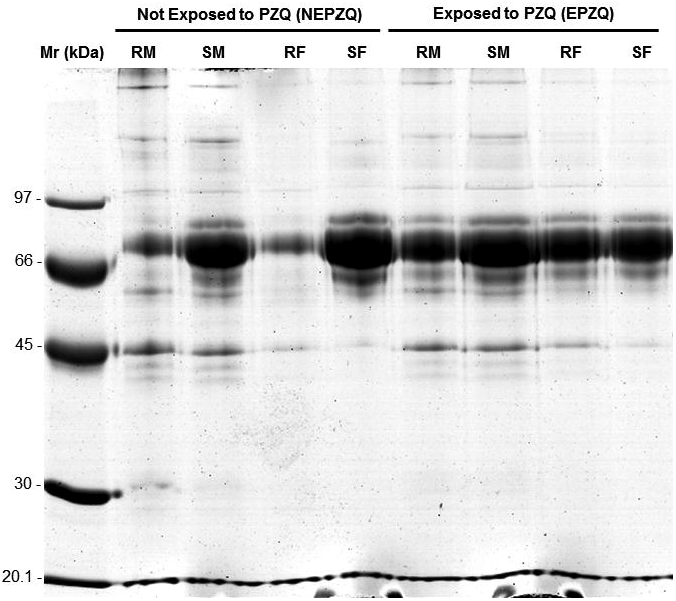
SDS-PAGE gel of the protein preparations, confirming the quality of the protein extracts studied. RM – Resistant males; SM – Susceptible males; RF – Resistant females; SF – Susceptible females; Mr – Molecular reference.

Analytical 2-DE gels were produced using 13 cm, pH 3-10NL strips and SDS-PAGE 12%, stained by Coomassie Blue to reproducibly resolve protein spots in a broad pH range and molecular weight, and posteriorly compare the protein pattern of *S. mansoni* proteome from the two strains (PZQ-resistant and PZQ-susceptible) not exposed (Figure 2) and exposed to PZQ (Figure 3).

**Figure 2.**
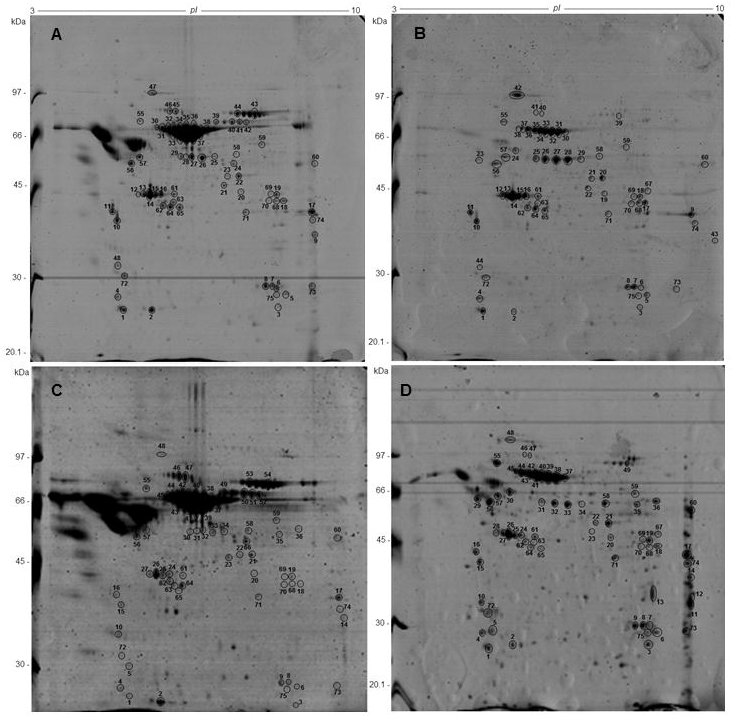
Two-dimensional gel electrophoresis of protein samples from *S. mansoni* adult worms not exposed to PZQ using 13 cm, pH 3-10NL strips and SDS-PAGE 12%, stained by Coomassie Blue. **(A)** SM-NEPZQ; **(B)** RM-NEPZQ; **(C)** SF-NEPZQ; **(D)** RF-NEPZQ. Numbers identify the spots, which were analyzed and identified by MS. All the identified proteins are listed in Supplementary material 1. The figure shows one representative experiment of three replicates.

**Figure 3.**
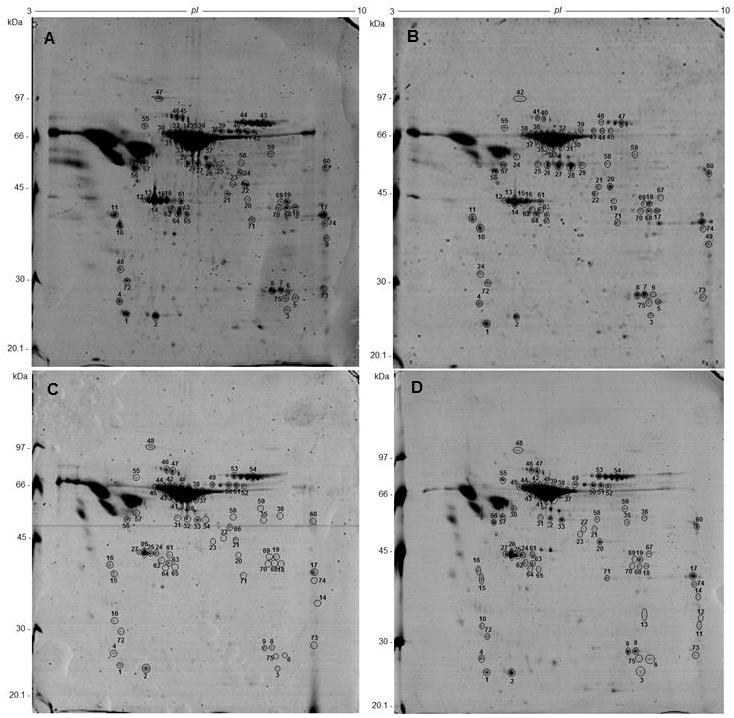
Two-dimensional gel electrophoresis of protein samples from *S. mansoni* adult worms exposed to PZQ using 13 cm, pH 3-10NL strips and SDS-PAGE 12%, stained by Coomassie Blue. **(A)** SM-EPZQ; **(B)** RM-EPZQ; **(C)** SF-EPZQ; **(D)** RF-EPZQ. Numbers identify the spots, which were analyzed and identified by MS. All the identified proteins are listed in Supplementary material 1. The figure shows one representative experiment of three replicates.

2-DE maps constructed with Coomassie Blue-stained gels showed reasonably comparable numbers of spots in all the samples. In total 133 ± 14, 265 ± 20, 142 ± 8, and 188 ± 34 spots were detected in proteins from RM-NEPZQ, RF-NEPZQ, SM-NEPZQ, and SF-NEPZQ, respectively (Table 1). For parasites exposed to PZQ, 203 ± 14, 133 ± 9, 220 ± 34 and 99 ± 19 spots were detected in RM-EPZQ, RF-EPZQ, SM-EPZQ and SF-EPZQ, respectively (Table 1). It is worth noting that 2-DE gels from RF-NEPZQ, RM-EPZQ and SM-EPZQ contain larger numbers of protein spots compared to other samples.

**Table 1.**
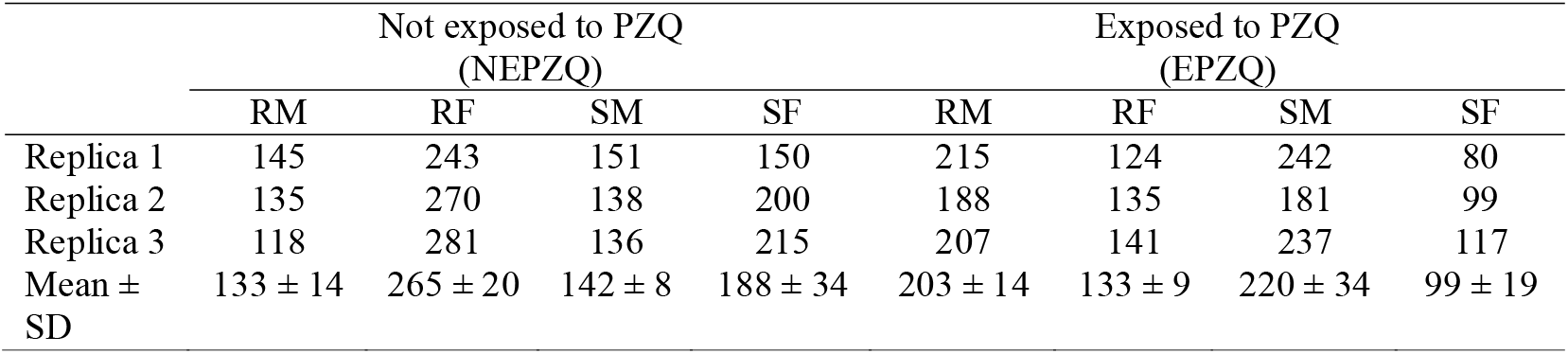
Summary comparison of the number of protein spots in the 2-DE maps for the eight different protein samples analyzed.

|  | Not exposed to PZQ<br>(NEPZQ) |  |  |  | Exposed to PZQ<br>(EPZQ) |  |  |  |
| --- | --- | --- | --- | --- | --- | --- | --- | --- |
|  | RM | RF | SM | SF | RM | RF | SM | SF |
| Replica 1 | 145 | 243 | 151 | 150 | 215 | 124 | 242 | 80 |
| Replica 2 | 135 | 270 | 138 | 200 | 188 | 135 | 181 | 99 |
| Replica 3 | 118 | 281 | 136 | 215 | 207 | 141 | 237 | 117 |
| Mean ± SD | 133 ± 14 | 265 ± 20 | 142 ± 8 | 188 ± 34 | 203 ± 14 | 133 ± 9 | 220 ± 34 | 99 ± 19 |

### 3.2 LC-MS/MS analysis and protein identification

The spots differentially expressed were excised from preparative gels of each sample, digested by trypsin and identified by LC-MS/MS. For RM-NEPZQ, 64 spots were successfully analyzed by LC-MS/MS, as well as 69 from RF-NEPZQ, 67 from SM-NEPZQ, 68 from SF-NEPZQ, 68 from RM-EPZQ, 69 from RF-EPZQ, 67 from SM-EPZQ, and 66 spots from SF-EPZQ (Supplementary material 1). The MS/MS results were employed to search the database (NCBInr) by Mascot search engine, and the matched proteins are listed in Supplementary material 1 and 2. Some proteins were identified in only one individual spot, but on several occasions, more than one spot in a gel corresponded to the same protein or better, same protein species (Figure 2, Figure 3 and Supplementary material 1).

Sixty individual protein species were identified on samples from parasites not exposed to PZQ, of which 45 were present in RM-NEPZQ, 52 in RF-NEPZQ, 45 in SM-NEPZQ and 47 proteins in SF-NEPZQ (Supplementary material 1). In this group of NEPZQ parasites, 35 proteins were common to all the four protein extracts, eight occurred only in resistant worms, eight only in susceptible worms, four in female parasites, and three were only present in resistant females. Interestingly, two proteins that have shown to be common in RM-NEPZQ, RF-NEPZQ and SM-NEPZQ preparations, namely serine/threonine phosphatase and troponin I, did not appear in SF-NEPZQ (Figure 4, Figure 5A and Table 2).

**Figure 4.**
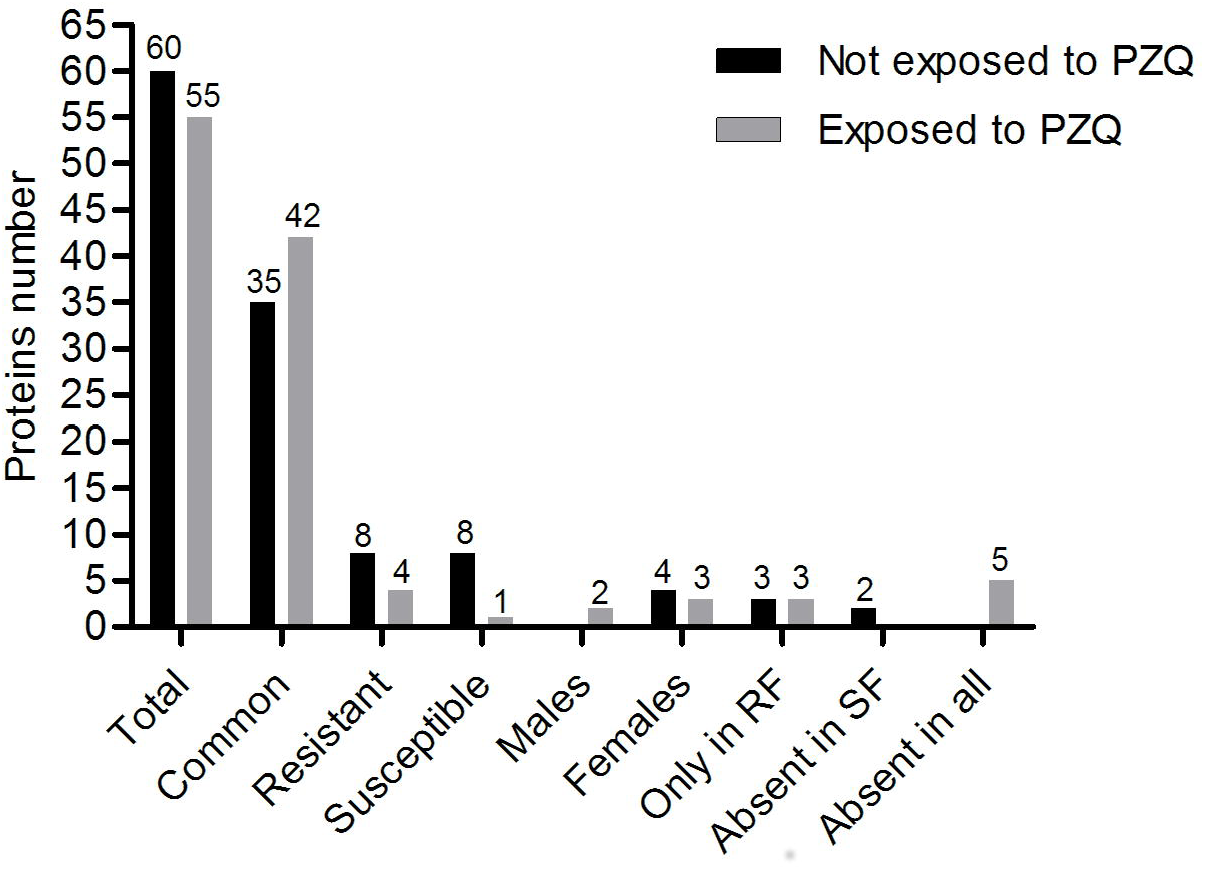
Number of shared proteins identified between and among the protein preparations from parasites not exposed and exposed to PZQ.

**Figure 5.**
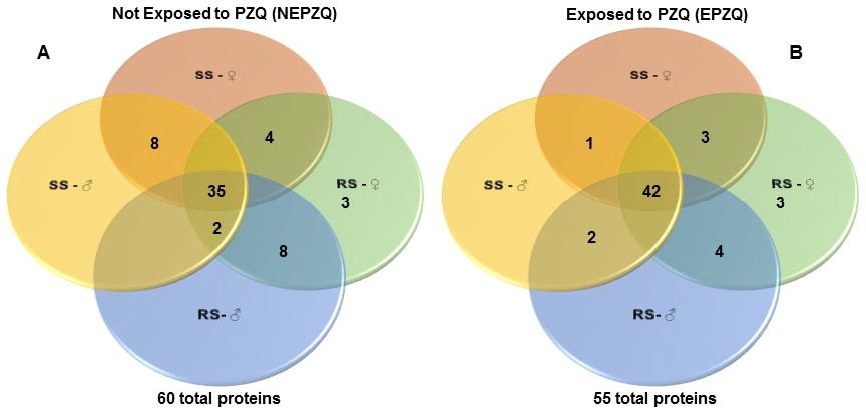
Venn diagram showing the number of shared proteins identified between and among the protein preparations from parasites. **(A)** Not exposed to PZQ (NEPZQ); **(B)** Exposed to PZQ. RS-♂: resistant strain males; RS-♀: resistant strain females; SS-♂: susceptible strain males; SS-♀: susceptible strain females. Common spots identified between and among the samples are represented overlapped by the circles.

**Table 2.**
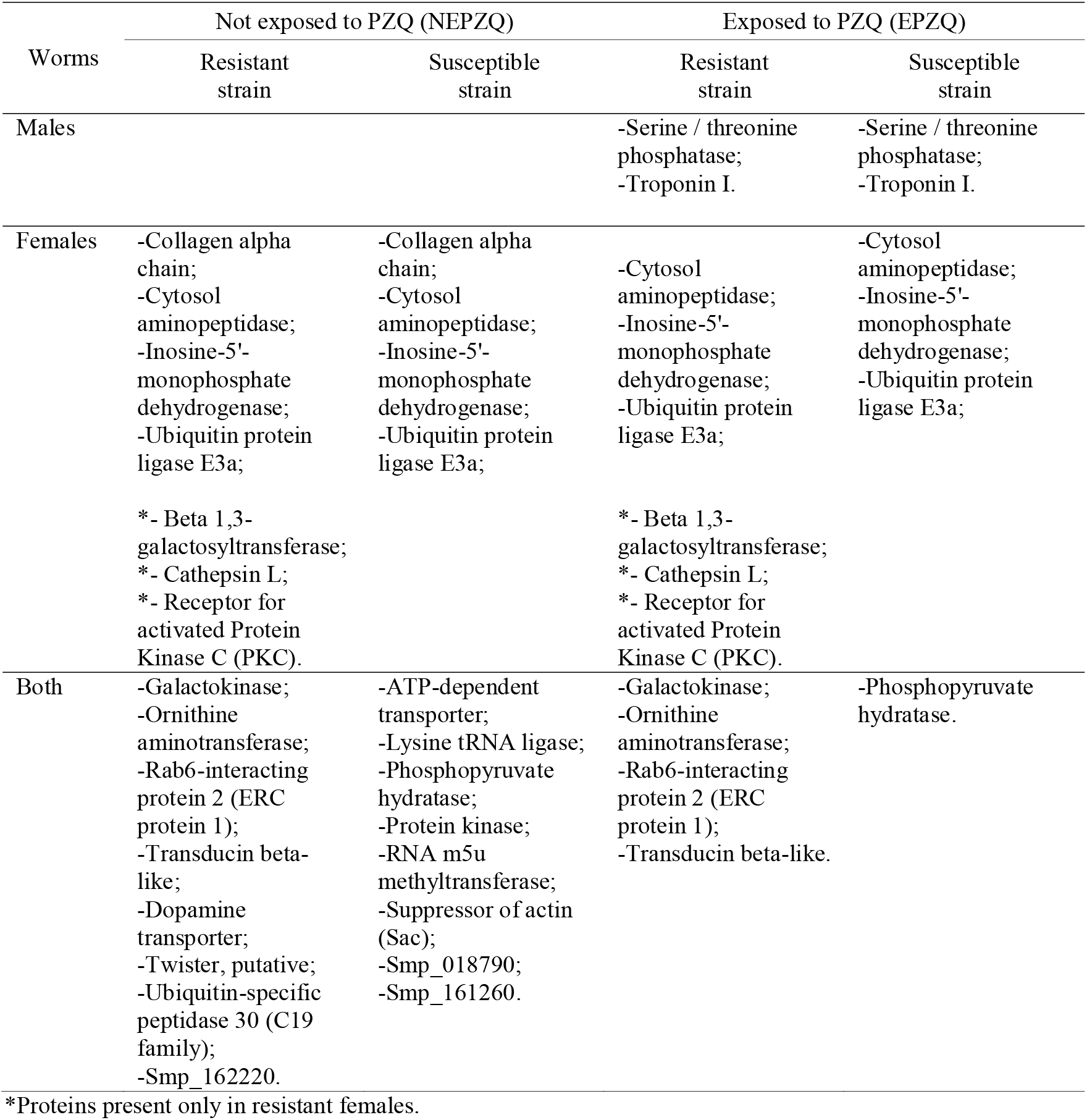
Specific proteins identified in each group analyzed.

The total number of proteins identified was reduced to 55 proteins on the protein extracts from parasites exposed to PZQ. Forty-eight of those proteins were present in RM-EPZQ, 52 in RF-EPZQ, 45 in SM-EPZQ, and 46 in SF-EPZQ (Supplementary material 1). Forty-two proteins appeared to be common to all the four proteins preparations of EPZQ parasites. In addition, four proteins were present only in resistant strain, one in susceptible strain, three were exclusive of female parasites, two proteins were present only in male parasites and three were only in resistant females. Five proteins that were present on parasites not exposed to PZQ, namely, collagen alpha chain, dopamine transporter, twister (putative), ubiquitin-specific peptidase 30 (C19 family), and Smp_162220, did not appear here (Figure 4, Figure 5B and Table 2).

### 3.3 Molecular function of identified proteins

The proteins identified by MS/MS, and summarized in Supplementary material 1 and 2, were categorized by their molecular function, according to information obtained from the GO database (Table 3), as described in the Material and methods section. When proteins had another function annotation, they were shown in brackets. The biological process and subcellular localization assigned to each protein in that database are also included in the Table 3. The results allowed the identification of proteins categorized as binding, catalytic, transport, regulation of muscle contraction, chaperone, motor, structural activities and proteins of unknown functions. Among the molecules identified as binding proteins, most of them were ATP, nucleotide, protein and ion binding proteins. The proteins categorized correspond to a variety of biological processes, nevertheless most of them were glycolytic enzymes and proteins related to metabolic process.

**Table 3.**
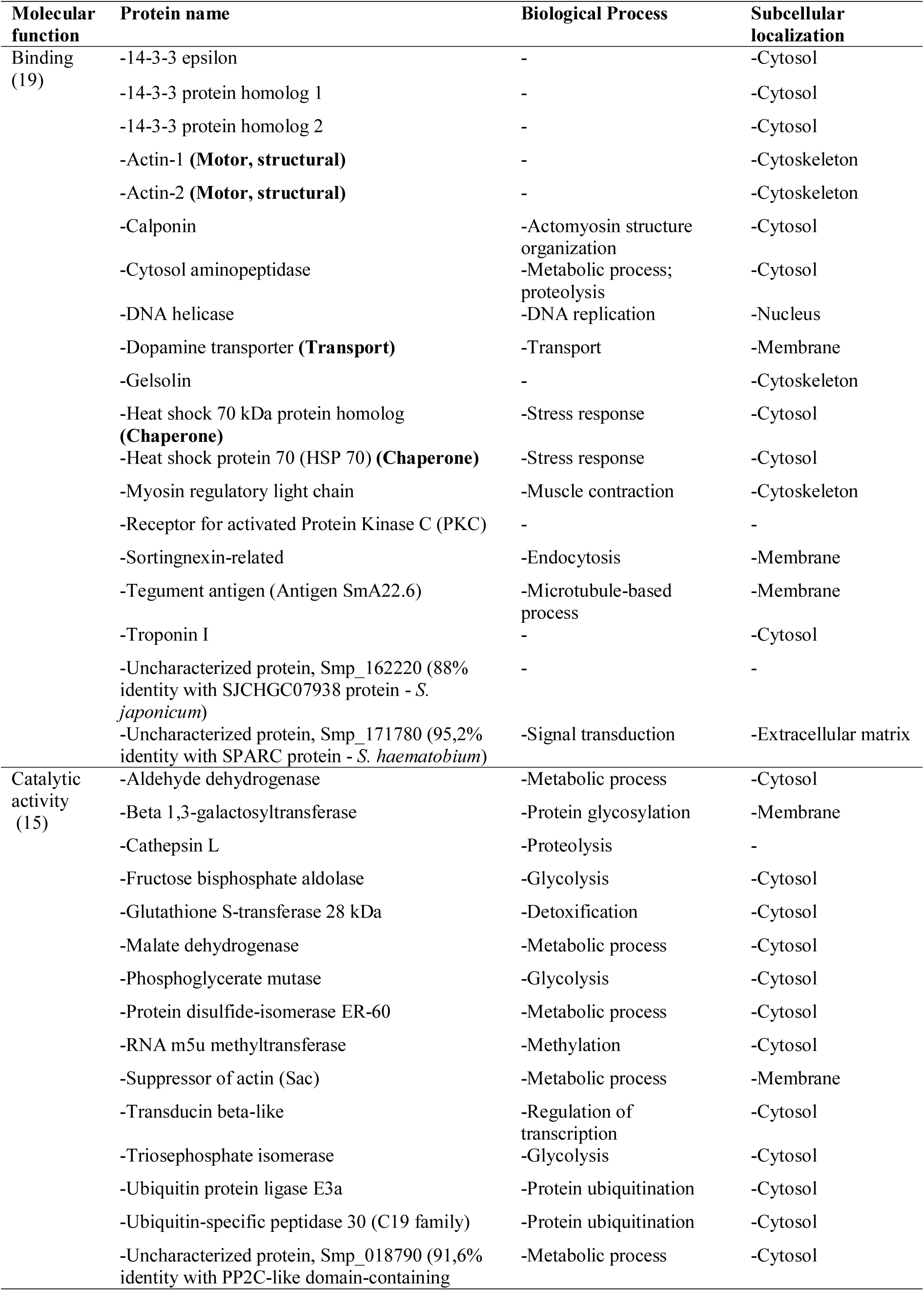
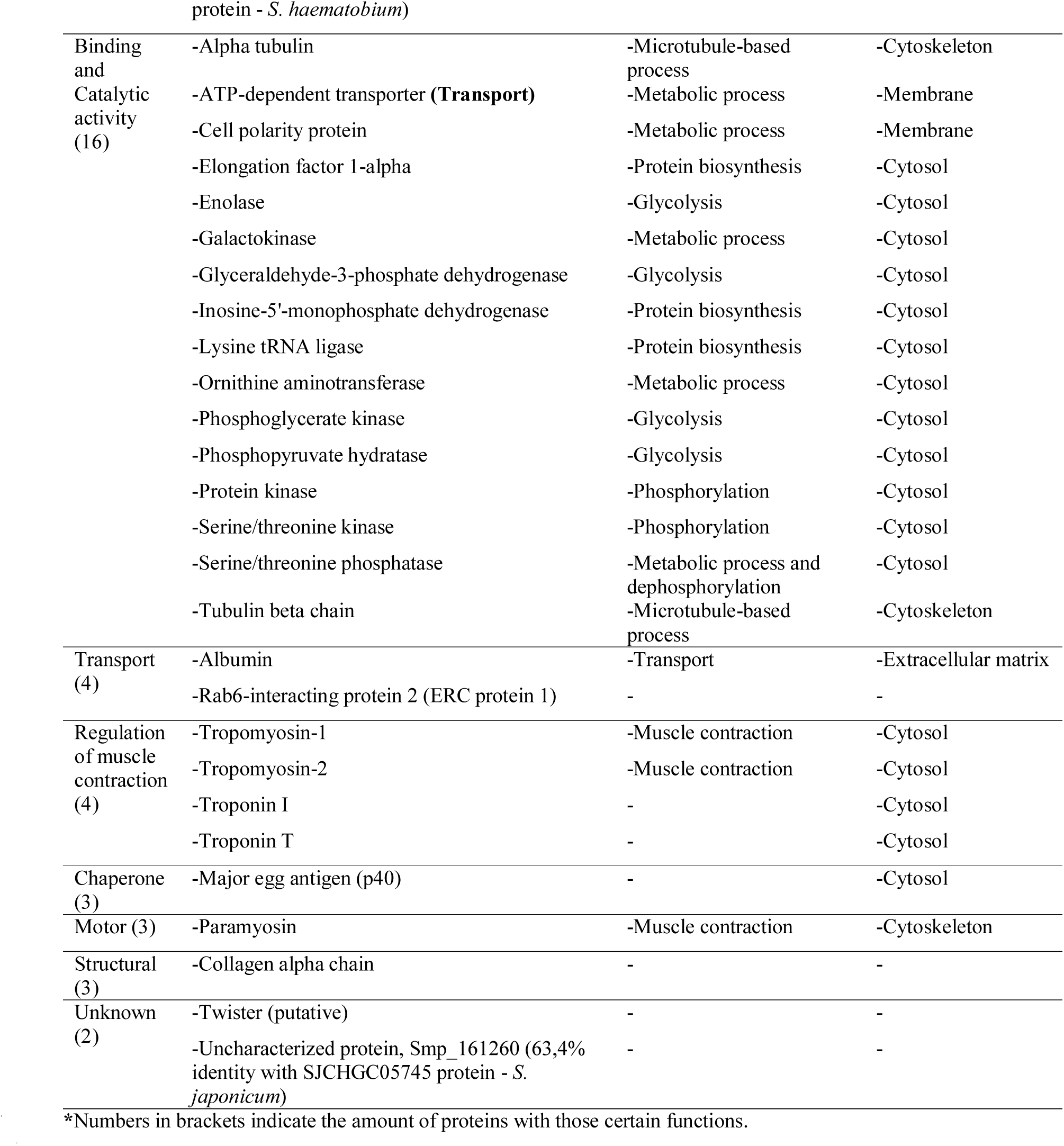
Proteins identified by their MS/MS and categorized by their molecular function according to information obtained from GO database.

Regarding to the subcellular localization, the proteins identified were classified, as cytoskeletal, cytosolic, nuclear, membrane proteins and some of them were located on extracellular matrix. Among them, those most abundantly identified were cytosolic proteins. There were fifteen and seven proteins whose biological process and subcellular localization, respectively, were not predicted (Table 3).

## 4. Discussion

Nowadays the control of schistosomiasis is based on PZQ, the only drug available for its treatment, which heavily relies on massive chemotherapy (Doenhoff et al., 2000; Cioli and Pica-Mattoccia, 2003; Hagan et al., 2004). However, the report of PZQ-resistance cases by *S. mansoni* has become a serious problem that needs to be solved. Several reports from our group and others have suggested that resistance to *Schistosoma* infection can be acquired naturally or induced by drug (Kabatereine et al., 1999; Corrêa-Oliveira et al., 2000; Black et al., 2010; Pinto-Almeida et al., 2015). Besides experimental evidences, reports of treatment failure in Senegal and Egypt in isolates with reduced susceptibility to PZQ were obtained (Stelma et al., 1995; Ismail et al., 1996) and further *in vitro* experiments have confirmed the development of PZQ-resistance (Fallon and Doenhoff, 1994; Ismail et al., 1999; Liang et al., 2001; Liang et al., 2010; Pinto-Almeida et al., 2015). Our group in particular has shown that resistance may be developed by drug pressure and as discussed in the Introduction section, we developed an important model that allows the laboratory study of PZQ-resistance in *S. mansoni* (Pinto-Almeida et al., 2015). Taking advantage of our drug resistant strain, the present study represents the first report of a *S. mansoni* PZQ-resistant strain proteomic analysis, comparing this strain with its isogenic counterpart susceptible strain, which is different only for the genetic determinants accounting for the PZQ-resistance phenotype.

In the present study we identified 60 different proteins on *S. mansoni* proteome. All those proteins were present in worms not exposed to PZQ, but some of them disappeared when these worms were exposed to the same drug. This result could possibly indicate an effect of PZQ exposure on protein expression in resistant and susceptible strains. Although previous studies of *Schistosoma* proteome had been performed using protein extracts and *Schistosoma* species different from ours, some proteins, such as 14-3-3 protein, HSP-70, GAPDH, glutathione S-transferase 28 kDa, enolase, fructose-bisphosphate aldolase, actin, triose phosphate isomerase, calponin, elongation factor 1-α, phosphoglycerate kinase, phosphoglycerate mutase, myosin, and paramyosin were commonly identified (Curwen et al., 2004; Mutapi et al., 2005; Braschi et al., 2006; Guillou et al., 2007; Pérez-Sánchez et al., 2008; Zhong et al., 2010; Boukli et al., 2011; Ludolf et al., 2014). In addition, some proteins that have already been tested as vaccine candidates, as glutathione S-transferase 28 kDa (Pearce, 2003), triose phosphate isomerase (Reynolds et al., 1994; Pearce, 2003) and paramyosin (Pearce, 2003) were also identified in the present study. Major egg antigen, troponin T, disulphide-isomerase ER-60 and actin, proteins that we also found, have already been clustered as immunoreactive proteins in serum pools of infected or non-infected individuals from endemic area (Ludlf et al., 2014).

Looking at the proteomes from both genders, in this survey, four proteins were only expressed in females from both strains, even under exposure to PZQ, namely, cytosol aminopeptidase, inosine-5’-monophosphate dehydrogenase (IMPDH), ubiquitin protein ligase E3a, and collagen alpha chain. Cytosol/leucine aminopeptidase catalyzes the hydrolysis of amino-acid residues from N-terminus of proteins and peptides (Piacenza et al., 1999), and it has already been assessed as a vaccine candidate against the infection of *Fasciola hepatica* (Acosta et al., 2008). This protein has previously been identified in *Schistosoma mansoni* eggs (Rinaldi et al., 2009). Regarding IMPDH, this protein is responsible for the rate-limiting step in guanine nucleotide biosynthesis (Prosise and Luecke, 2003), and it has previously been identified in *Schistosoma* genome and transcriptome (Protasio et al., 2012). E3 ligase enzyme catalyzes protein ubiquitination, which regulates various biological processes through covalent modification of proteins and transcription factors, and ubiquitin is the most important protein of this process (Sun and Chen, 2004; Santos et al., 2007). It has been suggested that ubiquitination is of interest in *S. mansoni* because this process could be a potential target for the design of new drugs (Guerra-Sá et al., 2005), being ubiquitin protein ligase E3a a good target to be studied. In regard to collagen alpha chain, Yang and colleagues (Yang et al., 2012)described that silencing the expression of a type of collagen (type V collagen) significantly affects the spawning and egg hatching of *S. japonicum*, and it also affects the morphology of the worms (Yang et al., 2012). Therefore, it would be very interesting evaluate the role of each of these proteins in PZQ-resistance, specially, collagen alpha chain, since it seems to occur morphological alterations in eggs and worms of *S. mansoni* PZQ-resistant strain (Pinto-Almeida et al., 2016). Moreover, females of this species, the only gender in which those proteins were found in this work, are more tolerant to PZQ treatment than males (Pinto-Almeida et al., 2015).

Another large difference between the proteome of both genders was the expression of troponin I and serine/threonine-protein phosphatase. Those proteins were present in males independently of drug exposure, but in females they were only present in resistant females not exposed to PZQ. Troponin I belongs to the troponin complex that mediate Ca^2+^-regulation that governs the actin-activated myosin motor function in striated muscle contraction (Wei and Jin, 2011). On the other side, protein kinases and phosphatases, as is the case of serine/threonine-protein phosphatase, are essential for normal functioning of signaling pathways since, it is well known that reversible phosphorylation of proteins is a ubiquitous mechanism crucial for regulation of most cellular functions (Luan, 2003). In *S. mansoni*, a phosphatase 2B (calcineurin) has been described as a heterodimer with a catalytic subunit and a regulatory subunit, which bind to Ca^2+^ increased the phosphatase activity (Mecozzi et al., 2000). Thus, protein phosphatases represent crucial molecules for the parasite and hence potential chemotherapeutic targets (Daher et al., 2006). Those differences in proteomes of males and females of *S. mansoni* represents a remarkable finding that is in agreement with other works (Cheng et al., 2005; Pérez-Sánchez et al., 2008), who have described a differential protein expression between males and females of *Schistosoma* spp.. However, these proteins are different from those described here, but it should be taken into account that those studies were performed with different species of *Schistosoma* (Cheng et al., 2005; Pérez-Sánchez et al., 2008).

Regarding to resistant strain parasites, it is notable the finding of eight proteins exclusively expressed in those *S. mansoni* worms. From those eight proteins, dopamine transporter, twister (putative), ubiquitin-specific peptidase 30 (C19 family), and uncharacterized protein smp_162220 are not present in *S. mansoni* PZQ-exposed worms. However, galactokinase, ornithine aminotransferase, Rab6-interacting protein 2 (ERC protein 1) and transducin beta-like remained after drug exposure. Dopamine/norepinephrine transporter (*SmDAT*) gene transcript, characterized in *S. mansoni*, is essential for the survival of the parasite as it causes muscular relaxation and a lengthening in the parasite, controlling movement (Larsen et al., 2011). Galactokinase catalyzes the second step of the Leloir pathway, a metabolic pathway found in most organisms for the catabolism of β-D-galactose to glucose 1-phosphate (Frey, 1996). Galactokinase and hexokinase have similar enzymatic function on sugar phosphorylation (Bork et al., 1993), and characterization of schistosome hexokinase has been described as pertinent to understanding the metabolic response of *S. mansoni* cercariae to an increased glucose availability (Tielens et al., 1994). Ornithine aminotransferase was already identified in *S. mansoni* (Roger et al., 2008) and it has been characterized as playing a central role in ornithine biosynthesis (Gafan et al., 2001). It seems responsible for catalyzing the transfer of the delta-amino group of L-ornithine to 2-oxoglutarate, producing L-glutamate-gamma-semialdehyde, that in turn spontaneously cyclizes to pyrroline-5- carboxylate, and L-glutamate (Haslett et al., 2004). Rab6-interacting protein 2 is a member of a family of RIM-binding proteins, which are presynaptic active zone proteins that regulate neurotransmitter release (Wang et al., 2002). Ubiquitin-specific peptidase 30 (C19 family) belongs to a metabolic pathway that had previously been associated to development of artemisinin and artesunate-resistance in *Plasmodium chabaudi* (Hunt et al., 2007), which is a very interesting result. All those proteins specifically found in the resistant strain should be further studied to better understand if they could possibly have a fundamental role in PZQ-resistance development.

Yet, for the resistant strain parasites, there are three proteins, beta 1,3-galactosyltransferase, cathepsin L, and receptor for activated Protein Kinase C (PKC) that are exclusive to resistant females, even after exposure to PZQ. Beta 1,3-galactosyltransferase has previously been identified in *Schistosoma* genome and transcriptome (Protasio et al., 2012). Cathepsin L activity is believed to be involved in hemoglobin digestion by adult schistosomes (Dalton et al., 1996), and these authors suggested the involvement of cathepsin proteinases in several key functions render them as potential targets to novel antiparasitic chemotherapy and immunoprophylaxis. Putative PKC exists in kinomes of the blood flukes *S. mansoni* (Berriman et al., 2009; Andrade et al., 2011), *S. japonicum* (Zhou et al., 2009) and *S. haematobium* (Young et al., 2012), and regulates movement, attachment, pairing, and egg release in *S. mansoni*, being considered a potential target for chemotherapeutic treatment against schistosomiasis (Ressurreição et al., 2014). These results really suggest a possible relationship between those proteins and PZQ-resistance in *S. mansoni* females, possibly being responsible for the exacerbated resistance demonstrated by those females in previous work (Pinto-Almeida et al., 2015). Concerning the PZQ-susceptible strain, it was noted eight proteins that only appeared in this strain. Phosphopyruvate hydratase (enolase), an important glycolytic enzyme that has the functions of activating the plasminogen, involving in the processes of infection and migration of parasites, reducing the immune function of the host as well as preventing parasites from the immune attack of the host (Gao and Yu, 2014), is the only protein from those eight proteins that continued to be expressed after PZQ exposure. This possibly suggests a relationship of this protein with the PZQ-susceptibility/resistance. However, more studies are necessary to investigate this hypothesis.

All those results together represent an important finding for the study of PZQ-resistance/susceptibility in *S. mansoni*, since they allow comparing directly the proteome under both conditions. We believe that the most promising candidates are proteins that appeared associated only to one of the strains, especially those with functions possibly related with the phenotypic alterations observed in (Pinto-Almeida et al., 2016), or previously associated with resistance by others parasites to different drugs. These candidates require special attention in more studies, assessing for instance the level of protection induced by these proteins in animal models infected by both *S. mansoni* strains, as they may have some involvement on PZQ-resistance phenomenon.

Here, we studied for the first time the proteome of a PZQ-resistant *S. mansoni* strain and the respective isogenic susceptible strain, a very important step towards understanding PZQ-resistance. This allowed us to identify proteins possibly associated with PZQ-resistance. These are proteins that were found to be differentially expressed between the strains. Since these proteins may be related with the PZQ-resistance phenomenon their functional characterization should pursue in future studies aimed at identifying new drug targets for schistosomiasis control. The identification of the proteins putatively associated with PZQ-resistance in *S. mansoni* permits also to investigate the possibility of developing a diagnosis test to distinguish patients carrying PZQ-resistant strains from those with PZQ-susceptible *S. mansoni*. The development of such a test would constitute a major step towards schistosomiasis control as it would render possible to adjust drug administration in order to increase treatment efficacy, perhaps even by combining PZQ with efflux pumps inhibitors as suggested previously (Pinto-Almeida et al., 2015). The proteome analysis made it also possible to identify proteins that were present only in females, being these good targets to identify the mechanisms underlying the decreased PZQ-susceptibility of females, when compared to males. Furthermore, some of these proteins may constitute targets, for schistosomiasis control. They should, therefore, be object of further analysis. In this context, the development and use of other techniques, such as genetic manipulation methods, will be crucial to further unravel the phenomenon/problem of PZQ-resistance/tolerance/loss of susceptibility. Therefore, this work opens doors to other PZQ-resistance studies, and could possibly represent a basis to find a solution to the PZQ-resistance problem in a disease that affects millions of people worldwide.

## Abbreviations

2-DE: two-dimensional electrophoresis
ACN: acetonitrile
BH: Belo Horizonte
DHPC: diheptanoyl-snglycero-3-phosphatidylcholine
DTT: dithiothreitol
EPZQ: exposed to Praziquantel
FA: formic acid
GO: gene ontology
IAA: iodoacetamide
IEF: isoelectric focusing
IHMT/UNL: Instituto de Higiene e Medicina Tropical, Universidade Nova de Lisboa
IMPDH: Inosine-5’-monophosphate dehydrogenase
IPG: immobilized pH gradient
LC-MS/MS: liquid chromatography–tandem mass spectrometry
MS: mass spectrometry
NEPZQ: not exposed to Praziquantel
PKC: protein kinase C
PZQ: Praziquantel
RF: resistant female
RM: resistant male
RS: resistant strain
SDS-PAGE: sodium dodecyl sulfate – polyacrylamide gel electrophoresis
SF: susceptible female
SM: susceptible male
SS: susceptible strain
TFA: trifluoracetic acid

## Acknowledgments

We acknowledge the Mass Spectrometry Laboratory at Brazilian Biosciences National Laboratory, CNPEM, Campinas, Brazil for their support with the mass spectrometry analysis. The authors would like to thank Professor Ana Tomás from IBMC and GABBA program, Porto, Portugal for very helpful suggestions that improved greatly this paper.

## Supplementary materials

**Supplementary material 1.** Proteins and spots identified in the samples from parasites not exposed and exposed to PZQ.

**Supplementary material 2.** Data related with protein and peptide identification.

## References

Acosta, D., Cancela, M., Piacenza, L., Roche, L., Carmona, C., and Tort, J.F. (2008). *Fasciola hepatica* leucine aminopeptidase, a promising candidate for vaccination against ruminant fasciolosis. Mol. Biochem. Parasitol. 158, 52–64. doi: 10.1016/j.molbiopara.2007.11.011

Al-Sherbiny, M., Osman, A., Barakat, R., El Morshedy, H., Bergquist, R., and Olds, R. (2003). *In vitro* cellular and humoral responses to *Schistosoma mansoni* vaccine candidate antigens. Acta Trop. 88, 117–130.

Andrade, L.F., Nahum, L.A., Avelar, L.G., Silva, L.L., Zerlotini, A., Ruiz, J.C., et al. (2011). Eukaryotic protein kinases (ePKs) of the helminth parasite *Schistosoma mansoni*. BMC Genomics 12, 215. doi: 10.1186/1471-2164-12-215

Babu, G.J., Wheeler, D., Alzate, O., and Periasamy, M. (2004). Solubilization of membrane proteins for two-dimensional gel electrophoresis: identification of sarcoplasmic reticulum membrane proteins. Anal Biochem. 325, 121–125.

Berriman, M., Haas, B.J., LoVerde, P.T., Wilson, R.A., Dillon, G.P., Cerqueira, G.C., et al. (2009). The genome of the blood fluke *Schistosoma mansoni*. Nature 460, 352–358. doi: 10.1038/nature08160

Black, C.L., Mwinzi, P.N., Muok, E.M., Abudho, B., Fitzsimmons, C.M., Dunne, D.W., et al. (2010). Influence of exposure history on the immunology and development of resistance to human schistosomiasis mansoni. PloS Negl Trop. Dis. 4:e637. doi: 10.1371/journal.pntd.0000637

Bork, P., Sander, C., and Valencia, A. (1993). Convergent evolution of similar enzymatic function on different protein folds: the hexokinase, ribokinase, and galactokinase families of sugar kinases. Protein Sci. 2, 31–40.

Boukli, N.M., Delgado, B., Ricaurte, M., and Espino, A.M. (2011). *Fasciola hepatica* and *Schistosoma mansoni*: identification of common proteins by comparative proteomic analysis. J. Parasitol. 97, 852–861. doi: 10.1645/GE-2495.1

Bradford, M.M. (1976). A rapid and sensitive method for the quantitation of microgram quantities of protein utilizing the principle of protein-dye binding. Anal Biochem. 72, 248–254.

Braschi, S., Curwen, R.S., Ashton, P.D., Verjovski-Almeida, S., and Wilson, A. (2006). The tegument surface membranes of the human blood parasite *Schistosoma mansoni*: a proteomic analysis after differential extraction. Proteomics 6, 1471–1482.

Braschi, S., and Wilson, R.A. (2006). Proteins Exposed at the Adult Schistosome Surface Revealed by Biotinylation. Mol. Cell. Proteomics 5, 347–356.

Cass, C.L., Johnson, J.R., Califf, L.L., Xu, T., Hernandez, H.J., Stadecker, M.J., et al. (2007). Proteomic analysis of *Schistosoma mansoni* egg secretions. Mol. Biochem. Parasitol. 55, 84–93.

Cheng, G.F., Lin, J.J., Feng, X.G., Fu, Z.Q., Jin, Y.M., Yuan, C.X., et al. (2005). Proteomic analysis of differentially expressed proteins between the male and female worm of *Schistosoma japonicum* after pairing. Proteomics 5, 511–521.

Chitsulo, L., Engles, D., Montresor, A., and Savioli, L. (2000). The global status of schistosomiasis and its control. Acta Trop. 77, 41–51.

Cioli, D., and Pica-Mattoccia, L. (2003). Praziquantel. Parasitol. Res. 90, S3–S9.

Clauser, K.R., Baker, P., and Burlingame, A.L. (1999). Role of accurate mass measurement (+/- 10 ppm) in protein identification strategies employing MS or MS/MS and database searching. Anal Chem. 71, 2871–2882.

Corrêa-Oliveira, R., Caldas, I.R., and Gazzinelli, G. (2000). Natural versus drug-induced resistance in *Schistosoma mansoni* infection. Parasitol. Today 16, 397–399.

Crompton, D.W. (1999). How much human helminthiasis is there in the world? J. Parasitol. 85, 397–403.

Curwen, R.S., Ashton, P.D., Johnston, D.A., and Wilson, R.A. (2004). The *Schistosoma mansoni* soluble proteome: a comparison across four life-cycle stages. Mol. Biochem. Parasitol. 138, 57–66.

Daher, W., Cailliau, K., Takeda, K., Pierrot, C., Khayath, N., Dissous, C., et al. (2006). Characterization of *Schistosoma mansoni* Sds homologue, a leucine-rich repeat protein that interacts with protein phosphatase type 1 and interrupts a G2/M cell-cycle checkpoint. Biochem. J. 395, 433–441.

Dalton, J.P., Clough, K.A., Jones, M.K., and Brindley, P.J. (1996). Characterization of the cathepsin-like cysteine proteinases of *Schistosoma mansoni*. Infect. Immun. 64, 1328–1334.

DeMarco, R., and Verjovski-Almeida, S. (2009). Schistosomes--proteomics studies for potential novel vaccines and drug targets. Drug. Discov. Today 14, 472–478. doi: 10.1016/j.drudis.2009.01.011

Doenhoff, M., Kimani, G., and Cioli, D. (2000). Praziquantel and the control of schistosomiasis. Parasitol. Today 16, 364–366.

Dvorák, J., Mashiyama, S.T., Braschi, S., Sajid, M., Knudsen, G.M., Hansell, E., et al. (2008). Differential use of protease families for invasion by schistosome cercariae. Biochimie 90, 345–358.

Fallon, P.G., and Doenhoff, M.J. (1994). Drug-resistant schistosomiasisL: resistance to praziquantel and oxamniquine induced in *Schistosoma mansoni* in mice is drug specific. Am. J. Trop. Med. Hyg. 51, 83–88.

Fenwick, A., Savioli, L., Engels, D., Robert Bergquist, N., and Todd, M.H. (2003). Drugs for the control of parasitic diseases: current status and development in schistosomiasis. Trends Parasitol. 19, 509–515.

Ferreira, M.S., de Oliveira, D.N., de Oliveira, R.N., Allegretti, S.M., Vercesi, A.E., and Catharino, R.R. (2014). Mass spectrometry imaging: a new vision in differentiating *Schistosoma mansoni* strains. J. Mass. Spectrom. 49, 86–92. doi: 10.1002/jms.3308

Frey, P.A. (1996). The Leloir pathway: a mechanistic imperative for three enzymes to change the stereochemical configuration of a single carbon in galactose. FASEB J. 10, 461–470.

Gafan, C., Wilson, J., Berger, L.C., and Berger, B.J. (2001). Characterization of the ornithine aminotransferase from *Plasmodium falciparum*. Mol. Biochem. Parasitol. 118, 1–10.

Gao, H., and Yu, C.X. (2014). Enolase and parasitic infection. Zhongguo Xue Xi Chong Bing Fang Zhi Za Zhi 26, 445–448.

Groth, D., Lehrach, H., and Hennig, S. (2004). GOblet: a platform for Gene Ontology annotation of anonymous sequence data. Nucleic Acids Res. 32, W313–317.

Guerra-Sá, R., Castro-Borges, W., Evangelista, E.A., Kettelhut, I.C., and Rodrigues, V. (2005). *Schistosoma mansoni*: functional proteasomes are required for development in the vertebrate host. Exp. Parasitol. 109, 228–236.

Guillou, F., Roger, E.. Mone, Y., Rognon, A., Grunau, C., Théron, A., et al. (2007). Excretory-secretory proteome of larval *Schistosoma mansoni* and *Echinostoma caproni*, two parasites of *Biomphalaria glabrata*. Mol. Biochem. Parasitol. 155, 45–56.

Hagan, P., Appleton, C.C., Coles, G.C., Kusel, J.R., and Tchuem-Tchuenté, L.A. (2004). Schistosomiasis control: keep taking the tablets. Trends Parasitol. 20, 92–97.

Hansell, E., Braschi, S., Medzihradszky, K.F., Sajid, M., Debnath, M., Ingram, J., et al. (2008). Proteomic analysis of skin invasion by blood fluke larvae. PLoS Negl Trop. Dis. 2:e262. doi: 10.1371/journal.pntd.0000262

Harrop, R., Coulson, P.S., and Wilson, R.A. (1999). Characterization, cloning and immunogenicity of antigens released by lung-stage larvae of *Schistosoma mansoni*. Parasitology 118, 583–594.

Haslett, M.R., Pink, D., Walters, B., and Brosnan, M.E. (2004). Assay and subcellular localization of pyrroline-5-carboxylate dehydrogenase in rat liver. Biochim. Biophys Acta 1675, 81–86.

Hu, W., Brindley, P.J., McManus, D.P., Feng, Z., and Han, Z.G. (2004). Schistosome transcriptomes: new insights into the parasite and schistosomiasis. Trends Mol. Med. 10, 217–225.

Hunt, P., Afonso, A., Creasey, A., Culleton, R., Sidhu, A.B., Logan, J., et al. (2007). Gene encoding a deubiquitinating enzyme is mutated in artesunate- and chloroquine-resistant rodent malaria parasites. Mol. Microbiol. 65, 27–40.

Huntley, R.P., Sawford, T., Mutowo-Muellenet, P., Shypitsyna, A., Bonilla, C., Martin, M.J., et al. (2015). The GOA database: Gene Ontology annotation updates for 2015. Nucleic Acids Res. 43, D1057–63. doi: 10.1093/nar/gku1113

Ismail, M., Botros, S., Metwally, A., William, S., Farghally, A., Tao, L.F., et al. (1999). Resistance to praziquantel: direct evidence from *Schistosoma mansoni* isolated from Egyptian villagers. Am. J. Trop. Med. Hyg. 60, 932–935.

Ismail, M., Metwally, A., Farghaly, A., Bruce, J., Tao, L.F., and Bennett, J.L. (1996). Characterization of isolates of *Schistosoma mansoni* from Egyptian villagers that tolerate high doses of praziquantel. Am. J. Trop. Med. Hyg. 55, 214–218.

James, C.E., Hudson, A.L., and Davey, M.W. (2009). Drug resistance mechanisms in helminths: is it survival of the fittest? Trends Parasitol. 25, 328–335. doi: 10.1016/j.pt.2009.04.004

Kabatereine, N.B., Vennervald, J.B., Ouma, J.H., Kemijumbi, J., Butterworth, A.E., Dunne, D.W., et al. (1999). Adult resistance to schistosomiasis mansoni: age-dependence of re-infection remains constant in communities with diverse exposure patterns. Parasitology 118, 101–105.

Kamel, E.G., El-Emam, M.A., Mahmoud, S.S., Fouda, F.M., and Bayaumy, F.E. (2011). Parasitological and biochemical parameters in *Schistosoma mansoni*-infected mice treated with methanol extract from the plants *Chenopodium ambrosioides*, *Conyza dioscorides* and *Sesbania sesban*. Parasitol. Int. 60, 388–392. doi: 10.1016/j.parint.2011.06.016

Katz, N., and Coelho, P.M. (2008). Clinical therapy of schistosomiasis mansoni: the Brazilian contribution. Acta Trop. 108, 72–78. doi: 10.1016/j.actatropica.2008.05.006

King, C.H. (2009). Toward the elimination of schistosomiasis. N. Engl. J. Med. 360, 106–109. doi: 10.1056/NEJMp0808041

Knudsen, G.M., Medzihradszky, K.F., Lim, K.C., Hansell, E., and McKerrow, J.H. (2005). Proteomic analysis of *Schistosoma mansoni* cercarial secretions. Mol. Cell. Proteomics 4, 1862–1875.

Kumagai, T., Maruyama, H., Hato, M., Ohmae, H., Osada, Y., Kanazawa, T., et al. (2005). *Schistosoma japonicum*: localization of calpain in the penetration glands and secretions of cercariae. Exp. Parasitol. 109, 53–57.

Larsen, M.B., Fontana, A.C., Magalhães, L.G., Rodrigues, V., and Mortensen, O.V. (2011). A catecholamine transporter from the human parasite *Schistosoma mansoni* with low affinity for psychostimulants. Mol. Biochem. Parasitol. 177, 35–41. doi: 10.1016/j.molbiopara.2011.01.006

Lewis, F.A. (1998). “Schistosomiasis,” in Current protocols in immunology, eds J.E. Coligan, A.M. Kruisbeek, D.H. Margulies, E.M. Shevach, W. Strober, and R. Coico (Hoboken, NJ: Wiley Interscience), 19.1.1–19.1.28.

Liang, Y.S., Coles, G.C., Doenhoff, M.J., and Southgate, V.R. (2001). *In vitro* responses of praziquantel-resistant and –susceptible *Schistosoma mansoni* to praziquantel. Int. J. Parasitol. 31, 1227–1235.

Liang, Y.S., Wang, W., Dai, J.R., Li, H.J., Tao, Y.H., Zhang, J.F., et al. (2010). Susceptibility to praziquantel of male and female cercariae of praziquantel-resistant and susceptible isolates of *Schistosoma mansoni*. J. Helminthol. 84, 202–207. doi: 10.1017/S0022149X0999054X

Liu, F., Cui, S.J., Hu, W., Feng, Z., Wang, Z.Q., and Han, Z.G. (2009). Excretory/secretory proteome of the adult developmental stage of human blood fluke, *Schistosoma japonicum*. Mol. Cell. Proteomics 8, 1236–1251. doi: 10.1074/mcp.M800538-MCP200

Luan, S. (2003). Protein phosphatases in plants. Annu. Rev. Plant. Biol. 54, 63–92.

Ludolf, F., Patrocínio, P.R., Corrêa-Oliveira, R., Gazzinelli, A., Falcone, F.H., Teixeira-Ferreira, A., et al. (2014). Serological screening of the *Schistosoma mansoni* adult worm proteome. PLoS Negl Trop. Dis. 8:e2745. doi: 10.1371/journal.pntd.0002745

Mecozzi, B., Rossi, A., Lazzaretti, P., Kady, M., Kaiser, S., Valle, C., et al. (2000). Molecular cloning of *Schistosoma mansoni* calcineurin subunits and immunolocalization to the excretory system. Mol. Biochem. Parasitol. 110, 333–343.

Mutapi, F., Burchmore, R., Mduluza, T., Foucher, A., Harcus, Y., Nicoll, G., et al. (2005). Praziquantel treatment of individuals exposed to *Schistosoma haematobium* enhances serological recognition of defined parasite antigens. J. Infect. Dis. 192, 1108–1118.

Pearce, E.J. (2003). Progress towards a vaccine for schistosomiasis. Acta Trop. 86, 309–313.

Pérez-Sánchez, R., Valero, M.L., Ramajo-Hernández, A., Siles-Lucas, M., Ramajo-Martín, V., and Oleaga,A. (2008). A proteomic approach to the identification of tegumental proteins of male and female *Schistosoma bovis* worms. Mol. Biochem. Parasitol. 161, 112–123. doi: 10.1016/j.molbiopara.2008.06.011

Perkins, D.N., Pappin, D.J., Creasy, D.M., and Cottrell, J.S. (1999). Probability-based protein identification by searching sequence databases using mass spectrometry data. Electrophoresis 20, 3551–3567.

Piacenza, L., Acosta, D., Basmadjian, I., Dalton, J.P., and Carmona, C. (1999). Vaccination with cathepsin L proteinases and with leucine aminopeptidase induces high levels of protection against fascioliasis in sheep. Infect. Immun. 67, 1954–1961.

Pinto-Almeida, A., Mendes, T., Armada, A., Belo, S., Carrilho, E., Viveiros, M., et al. (2015). The Role of Efflux Pumps in *Schistosoma mansoni* Praziquantel Resistant Phenotype. PLoS ONE 10:e0140147. doi: 10.1371/journal.pone.0140147

Pinto-Almeida, A., Mendes, T., de Oliveira, R.N., Corrêa, S.A.P., Allegretti, S.M., Belo, S., et al. (2016). Morphological characteristics of *Schistosoma mansoni* PZQ-resistant and –susceptible strains are different in presence of Praziquantel. Front. Microbiol. 7:594. doi:10.3389/fmicb.2016.00594

Prosise, G.L., and Luecke, H. (2003). Crystal structures of Tritrichomonasfoetus inosine monophosphate dehydrogenase in complex with substrate, cofactor and analogs: a structural basis for the random-in ordered-out kinetic mechanism. J. Mol. Biol. 326, 517–527.

Protasio, A.V., Tsai, I.J., Babbage, A., Nichol, S., Hunt, M., Aslett, M.A., et al. (2012). A systematically improved high quality genome and transcriptome of the human blood fluke *Schistosoma mansoni*. PLoS Negl Trop. Dis. 6:e1455. doi: 10.1371/journal.pntd.0001455

Ressurreição, M., De Saram, P., Kirk, R.S., Rollinson, D., Emery, A.M., Page, N.M., et al. (2014). Protein kinase C and extracellular signal-regulated kinase regulate movement, attachment, pairing and egg release in *Schistosoma mansoni*. PLoS Negl Trop. Dis. 8:e2924. doi: 10.1371/journal.pntd.0002924

Reynolds, S.R., Dahl, C.E., and Harn, D.A. (1994). T and B epitope determination and analysis of multiple antigenic peptides for the *Schistosoma mansoni* experimental vaccine triose-phosphate isomerase. J. Immunol. 152, 193–200.

Rinaldi, G., Morales, M.E., Alrefaei, Y.N., Cancela, M., Castillo, E., Dalton, J.P., et al. (2009). RNA interference targeting leucine aminopeptidase blocks hatching of *Schistosoma mansoni* eggs. Mol. Biochem. Parasitol. 167, 118–126. doi: 10.1016/j.molbiopara.2009.05.002

Roger, E., Mitta, G., Moné, Y., Bouchut, A., Rognon, A., Grunau, C., et al. (2008). Molecular determinants of compatibility polymorphism in the *Biomphalaria glabrata*/*Schistosoma mansoni* model: new candidates identified by a global comparative proteomics approach. Mol. Biochem. Parasitol. 157, 205–216.

Santos, D.N., Aguiar, P.H., Lobo, F.P., Mourão, M.M., Tambor, J.H., Valadão, A.F., et al. (2007). *Schistosoma mansoni*: Heterologous complementation of a yeast null mutant by SmRbx, a protein similar to a RING box protein involved in ubiquitination. Exp. Parasitol. 116, 440–449.

Shevchenko, A., Tomas, H., Havlis, J., Olsen, J.V., and Mann, M. (2006). In-gel digestion for mass spectrometric characterization of proteins and proteomes. Nat. Protoc. 1, 2856–2860.

Smyth, D., McManus, D.P., Smout, M.J., Laha, T., Zhang, W., and Loukas, A. (2003). Isolation of cDNAs encoding secreted and transmembrane proteins from *Schistosoma mansoni* by a signal sequence trap method. Infect. Immun. 71, 2548–2554.

Steinmann, P., Keiser, J., Bos, R., Tanner, M., and Utzinger, J. (2006). Schistosomiasis and water resources development: systematic review, meta-analysis, and estimates of people at risk. Lancet. Infect. Dis. 6, 411–425.

Stelma, F.F., Talla, I., Sow, S., Kongs, A., Niang, M., Polman, K., et al. (1995). Efficacy and side effects of praziquantel in an epidemic focus of *Schistosoma mansoni*. Am. J. Trop. Med. Hyg. 53, 167–170.

Sun, L., and Chen, Z.J. (2004). The novel functions of ubiquitination in signaling. Curr. Opin. Cell. Biol. 16, 119–126.

Tielens, A.G., van den Heuvel, J.M., van Mazijk, H.J., Wilson, J.E., and Shoemaker, C.B. (1994). The 50-kDa glucose 6-phosphate-sensitive hexokinase of *Schistosoma mansoni*. J. Biol. Chem. 269, 24736–24741.

van Balkom, B.W., van Gestel, R.A., Brouwers, J.F., Krijgsveld, J., Tielens, A.G., Heck, A.J., et al. (2005). Mass spectrometric analysis of the *Schistosoma mansoni* tegumental sub-proteome. J. Proteome Res. 4, 958–966.

Verjovski-Almeida, S., Leite, L.C., Dias-Neto, E., Menck, C.F., and Wilson, R.A. (2004). Schistosome transcriptome: insights and perspectives for functional genomics. Trends Parasitol. 20, 304–308.

Wang, W., Wang, L., and Liang, Y.S. (2012). Susceptibility or resistance of praziquantel in human schistosomiasis: a review. Parasitol. Res. 111, 1871–1877. doi: 10.1007/s00436-012-3151-z

Wang, Y., Liu, X., Biederer, T., and Südhof, T.C. (2002). A family of RIM-binding proteins regulated by alternative splicing: Implications for the genesis of synaptic active zones. Proc. Natl. Acad. Sci. USA. 99, 14464–14469.

Wei, B., and Jin, J.P. (2011). Troponin T isoforms and posttranscriptional modifications: Evolution, regulation and function. Arch. Biochem. Biophys 505, 144–154. doi: 10.1016/j.abb.2010.10.013

World Health Organization (WHO) (2013). Schistosomiasis: Progress Report 2001–2011 and Strategic Plan 2012–2020. Geneva: World Health Organization Press.

Yang, Y., Jin, Y., Liu, P., Shi, Y., Cao, Y., Liu, J., et al. (2012). RNAi silencing of type V collagen in *Schistosoma japonicum* affects parasite morphology, spawning, and hatching. Parasitol. Res. 111, 1251–1257. doi: 10.1007/s00436-012-2959-x

Young, N.D., Jex, A.R., Li, B., Liu, S., Yang, L., Xiong, Z., et al. (2012). Whole-genome sequence of *Schistosoma haematobium*. Nature Genet. 44, 221–225. doi: 10.1038/ng.1065

Zhong, Z.R., Zhou, H.B., Li, X.Y., Luo, Q.L., Song, X.R., Wang, W., et al. (2010). Serological proteome-oriented screening and application of antigens for the diagnosis of Schistosomiasis japonica. Acta Trop. 116, 1–8. doi: 10.1016/j.actatropica.2010.04.014

Zhou, Y., Zheng, H., Chen, Y., Zhang, L., Wang, K., Guo, J., et al. (2009). The *Schistosoma japonicum* genome reveals features of host-parasite interplay. Nature 460, 345–351. doi: 10.1038/nature08140

